# The impact of intravenous dodecafluoropentane on a murine model of acute lung injury

**DOI:** 10.1101/2020.08.17.253658

**Authors:** Jarrod M Mosier, Saad Sammani, Carrie Kempf, Evan Unger, Joe GN Garcia

## Abstract

Acute hypoxemic respiratory failure presents therapeutic challenges due to ventilation/perfusion mismatch and shunt. The goal of management strategies is to improve arterial oxygenation, however each management strategy presents risk to patients from iatrogenic injury. Intravenous oxygen therapeutics present an appealing option to improve arterial oxygenation without these risks. We used a two-hit murine model of acute lung injury to evaluate the effect of intravenous dodecafluoropentane (NanO_2_) on oxygen saturation and bronchoalveolar lavage cell count and protein. Mice were given intratracheal lipopolysaccharide and 20 hours later were intubated and ventilated with high tidal volumes. NanO_2_ was given by bolus injection at the initiation of mechanical ventilation and again at 2 hours, while oxygen saturation was measured every 15 minutes. At the conclusion of the experiment (4 hours), a bronchoalveolar lavage was performed. There was no difference in mean O_2_ saturation at time zero, however the difference between the mean O_2_ saturation immediately prior to injection and the mean first O_2_ saturation after injection in the control saline group were 91% and 83%, mean difference −7.5%; whereas mean O_2_ saturation in the NanO_2_ treated group rose from 89% to 91%, mean difference +2.5%, net difference 10% [95% CI: 2.7,17.3], p=0.01). There was a statistically significant difference in cell count, but not protein, on the bronchoalveolar lavage analysis. These data show that NanO_2_ rapidly improves oxygen saturation in a two-hit model of acute lung injury, and shows potential as an intravenous oxygen therapeutic in the management in acute hypoxemic respiratory failure.

## Introduction

Acute hypoxemic respiratory failure is one of the most common reasons for admission in critically ill patients. The underlying ventilator/perfusion (V/Q) mismatch and intrapulmonary shunt presents increased risk to patients when using noninvasive ventilatory strategies, during preoxygenation for intubation, or during mechanical ventilation. Noninvasive strategies (1) suffer high failure rates, which imposes disproportionately higher mortality mediated, particularly with NIPPV, by patient-self-induced lung injury (P-SILI).(2, 3) Intubation in these patients comes with high risk of rapid desaturation, largely contributing to the increased cardiac arrest rate.(4-6) Mechanical ventilation carries the risk of ventilator-induced lung injury (VILI) from injurious settings required to maintain systemic oxygenation.(7)

Given these risks, novel approaches to improving arterial oxygenation in presence of increased V/Q mismatch and intrapulmonary shunt are desired. Venovenous extracorproreal membrane oxygenation (ECMO), for example, permits improved arterial oxygenation and allows for lung protective ventilation essentially by increasing the central venous oxygen saturation so the injured native lung is not required to do as much work. While ECMO can be a useful tool for patients with refractory hypoxemia and poor lung compliance, it is invasive, resource intensive, and lacks outcomes data prohibiting widespread use. Perfluorocarbons were recently attempted as liquid ventilation, and were abandoned for lack of efficacy and potential harm in clinical trials.(8) However, while intratracheal perfluorocarbons have shown harm, *intravenous* perfluorocarbons have potential as a small particle oxygen therapeutic.

Dodecafluoropentane emulsion [NanO_2_™, NuvOx Pharma (Tucson, AZ)] is one such promising perfluorocarbon. The molecular characteristics of NanO_2,_ comprised of 2% weight/volume dodecafluoropentane emulsion (DDFPe)(9) make it highly effective at oxygen delivery at extremely low doses,(10) and an ideal intravenous agent for O_2_ delivery. In this study, we aimed to evaluate the potential of NanO_2_ to improve oxygenation in conditions of low V/Q mismatch and high intrapulmonary shunt by using a preclinical model of acute lung injury and measuring the difference in mean oxygen saturation after injection of NanO_2_ or saline and differences in BAL cell count and protein.

## Materials and Methods

### Murine Model

We used an established two-hit mouse model of acute lung injury for this experiment.(11) C57BL/J6 male mice (n=12, 8-10 week old) were anesthetized with intraperitoneal ketamine (100 mg/kg) and xylazine (5 mg/kg), and then intubated with a 20-gauge intravenous catheter. Lipopolysaccharide (0.5 mg/kg) was then given by intratracheal injection, mice were extubated and allowed to recover for 20 hours. After the recovery period, the mice were resedated with ketamine and xylazine by intraperitoneal injection, with additional doses and bupremorphone (0.3mg/kg) given as needed for pain and to ensure deep sedation during the experiment, reintubated and placed on a mechanical ventilator (Harvard Apparatus, Boston, MA). Mice were ventilated to induce ventilator-induced lung injury by using a tidal volume of 30ml/kg, respiratory rate of 75 breaths/minute, and room air with no positive end-expiratory pressure for four hours.

The right internal jugular vein was surgically exposed and mice were injected with either NanO_2_ (0.6ml/kg) or an equivalent dose of saline (n=6/group) at time-zero (onset of mechanical ventilation) and again at two hours. Oxygen saturation was continuously monitored using the MouseStat® monitor (Kent Scientific Corporation, Torrington, CT) on the pad of a hind leg paw and measurements were recorded as an average over 30 seconds every 15 minutes.

At the termination of each experiment (after four hours of mechanical ventilation), bronchoalveolar lavage (BAL) fluid was collected by instilling 1 ml of HBSS (Invitrogen, Grand Island, NY) through the tracheal catheter, followed by slow recovery of the fluid. Cells were recovered from the resulting BAL fluid by centrifugation (500 *g*, 20 min, 4°C) and counted using an automated cell counter (TC20; Bio-Rad, Hercules, CA). All animal care procedures and experiments were approved by the University of Arizona Animal Care and Use Committee (Approval #13-490).

## Results

The mean oxygen saturation at the initiation of mechanical ventilation was 95%-96% for both the control and experiment groups. No significant differences in mean oxygen saturation were observed between groups for the first two hours, which both trended down to mean O_2_ saturations of 91% for the control group and 89% for the NanO_2_ treated group (**Fig. 1**). However, the mean O_2_ saturation differed significantly after the two-hour injection of NanO_2_ or saline. The difference between the mean O_2_ saturation immediately prior to injection and the mean first O_2_ saturation after injection in the control saline group were 91% and 83%, mean difference −7.5%; whereas mean O_2_ saturation in the NanO_2_ treated group rose from 89% to 91%, mean difference +2.5%, net difference 10% [95% CI: 2.7,17.3], p=0.01). O_2_ saturation continued to be higher for the mice receiving NanO_2_ for the remainder of the experiment (**Fig. 1**). BAL cell and protein values, reflecting lung inflammatory injury showed no difference in BAL protein content but a significant reduction in BAL cell count in mice receiving NanO_2_ vs controls (**Fig. 2**).

**Fig. 1:**
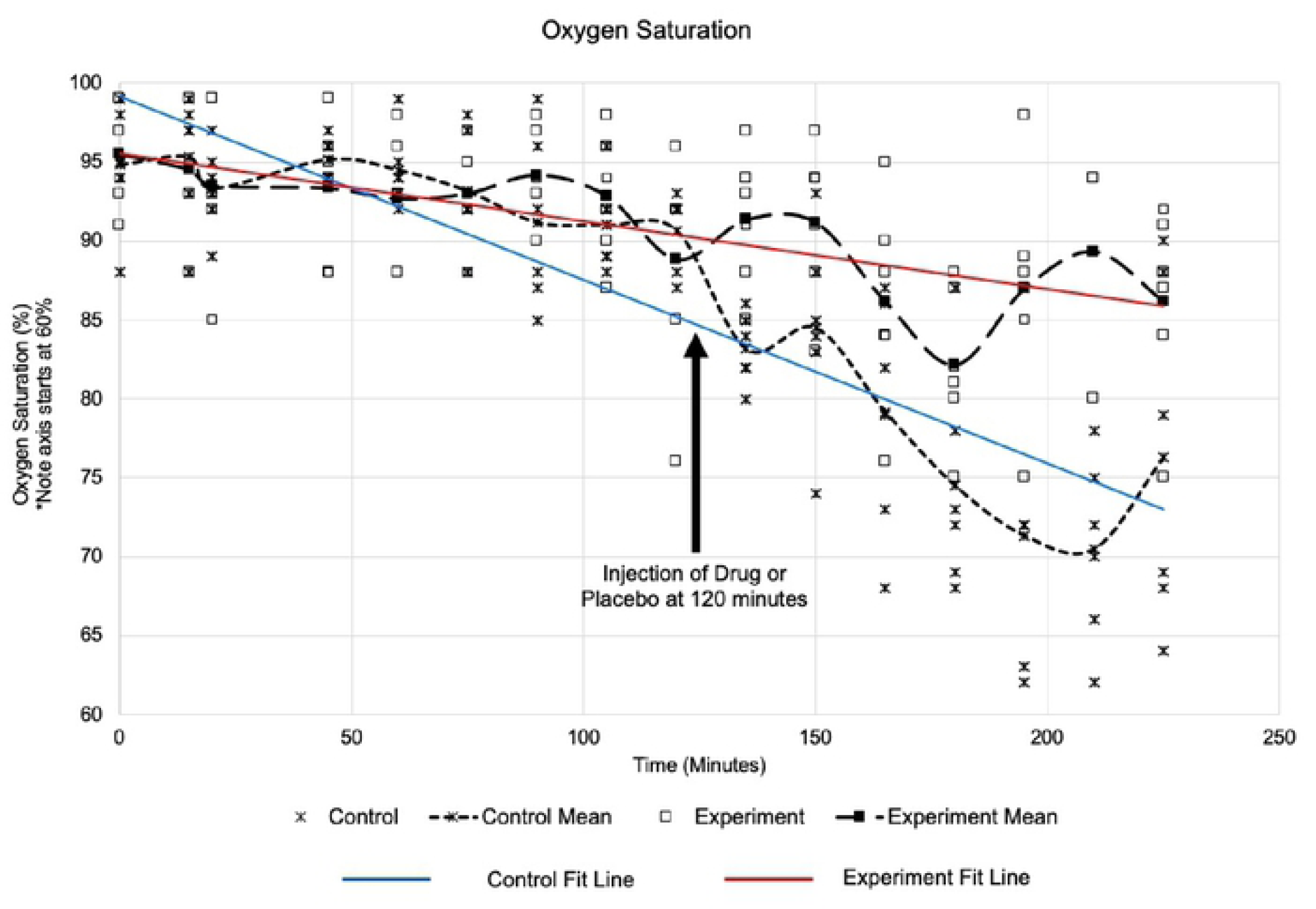
Oxygen saturations over time for control mice (⋇) and mice treated with NanO_2_ (open squares). Mean oxygen saturations for each group are represented by the dashed lines and the it lines for the control group (blue) and experiment mice (red) are also shown. There was no difference in mean oxygen saturation until the injection of drug or placebo at the two hour mark, at which point the NanO_2_ treated mice had improved oxygen saturations. Time zero is the initiation of mechanical ventilation with injurious tidal volumes, 20 hours after intratracheal lipopolysaccharide injection.

**Fig. 2:**
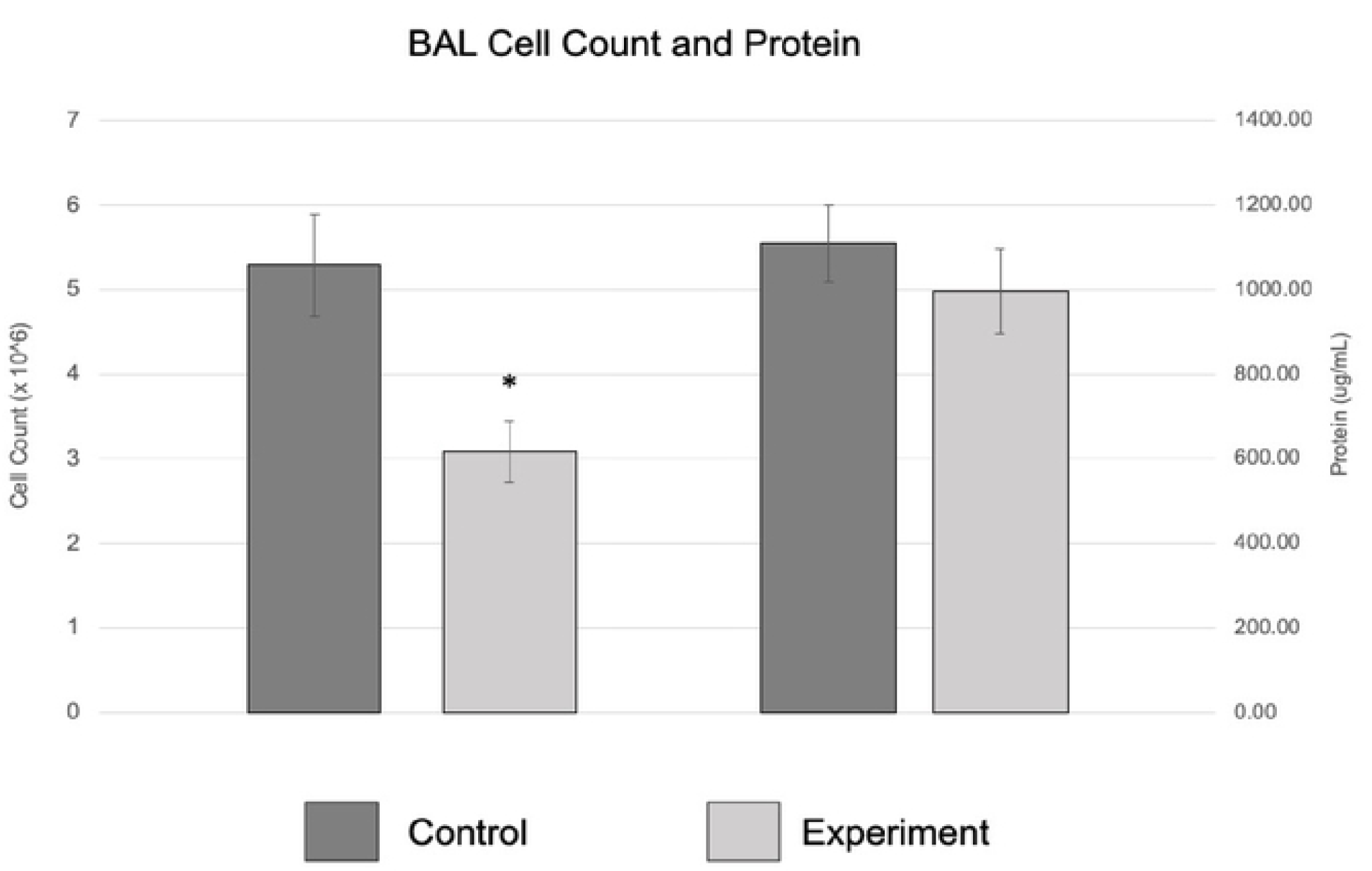
Bronchoalveolar lavage demonstrated a statistically significant decrease in cell count (p=0.01) with NanO_2_ treated mice, but no difference in protein content (p=0.33).

## Discussion

These results show that in a ‘two-hit’ model of acute lung injury, intravenously administered NanO_2_ rapidly improves oxygen saturation in hypoxemic mice within minutes of injection. The mice had similarly declining oxygen saturation trajectories until two hours, when the second injection of NanO_2_ rapidly increased oxygen saturation and changed the trajectory for the remainder of the experiment. Although the mechanism of action is not completely understood, NanO_2_ appears to improve arterial oxygen content despite disruptions in V/Q mismatch induced by both endothelial (LPS) and epithelial injury (volutrauma). It is interesting that a reduction in BAL cell count, but not BAL total protein was observed in NanO_2_-treated mice compared to controls. One hypothesis is that NanO2 may reduce oxidative stress leading to reduced cellular infiltration but not affecting capillary leak, but this mechanism is speculative and requires exploration.

NanO_2_ has demonstrated efficacy in preclinical models in reducing cerebral damage from acute ischemic stroke, (12-15) and decreasing myocardial damage in acute myocardial infarction.(16) NanO_2_ was also shown to rapidly improve PaO_2_ in a porcine model of intrapulmonary shunt induced by bead instillation into the bronchial tree,(17) as well as improve tissue hypoxia in preclinical hemorrhage, stroke, and acute chest syndrome models.(14, 18, 19) Additionally, NanO_2_ can be administered repeatedly to restore oxygenation,(17) which potential increases the therapeutic utility when treating acute hypoxemic respiratory failure. Our results add to the existing preclinical studies demonstrating the potential for NanO_2_ to restore systemic oxygenation and attenuate hypoxic tissue injury.

The improvement in oxygen saturation seen in this study is a potentially clinically translatable option for overcoming the challenges imposed by V/Q mismatch and shunt in patients with acute hypoxemic respiratory failure. This increase in oxygen saturation may augment conventional oxygen therapy or noninvasive respiratory strategies and prevent P-SILI. In patients requiring mechanical ventilation, the improved arterial oxygenation may allow for less aggressive ventilator settings thereby reducing ventilator-induced lung injury. Studies are needed in each of these patient populations but the potential is encouraging.

Patients with acute hypoxemic respiratory failure that do require intubation face a four-fold increase (adjusted odds) in risk of cardiac arrest with intubation compared to patients without hypoxemic respiratory failure.(4) Preoxygenation, which provides a reservoir of oxygen to resaturate hemoglobin during the apneic period that can last several minutes, is essential to avoiding critical desaturation leading to cardiac arrest.(20, 21) In patients with acute hypoxemic respiratory failure, particularly those with acute respiratory distress syndrome (ARDS), the efficacy of preoxygenation is limited by: a.) reduced functional residual capacity, b.) significant shunt physiology, and c.) a sudden worsening of V/Q mismatch when spontaneous breathing ceases after injection of induction medications.(5) These limitations worsen when advanced methods of preoxygenation are not used due to concern for viral aerosol production, such as with COVID-19 associated respiratory failure. The rapid increase in oxygen saturation seen in this experiment provides a potential solution to improving arterial oxygenation despite V/Q mismatch and shunt to augment the preoxygenation process, potentially attenuating intubation-related desaturation in critically ill patients and without producing aerosols.

Research is needed using various dosing strategies to evaluate the potential role of NanO_2_ for each of the therapeutic challenges mentioned in acute hypoxemic respiratory failure patients: noninvasive respiratory support, preoxygenation for intubation, and lung protective ventilation. Studies are also needed on the mechanism of action and safety in humans, however these results indicate that NanO_2_ is an appealing candidate for improving arterial oxygenation in the presence of V/Q mismatch and intrapulmonary shunt.

## Acknowledgements

This work was partially supported by an University of Arizona Health Sciences Career Development Award for Dr. Mosier.

